# Sex differences in the impact of nerve injury on Locus Coeruleus function and behaviour in mice

**DOI:** 10.1101/2024.11.06.622052

**Authors:** Patricia Mariscal, Adrián Martínez-Cortés, Meritxell Llorca-Torralba, Jone Razquin, Cristina Miguelez, Cristina Ulecia-Morón, Borja García-Bueno, Juan Carlos Leza, Lidia Bravo, Esther Berrocoso

## Abstract

Chronic pain often coexists with stress-related disorders such as anxiety and depression, with a higher prevalence in women. Rodent studies reveal significant sex differences in pain and stress-related behaviours, emphasising the need to explore underlying neurobiological mechanisms. The locus coeruleus (LC), the main noradrenergic nucleus in the brainstem, plays a crucial role in modulating pain and stress responses and exhibits sex-specific characteristics.

In this study, we investigated the effects of neuropathic pain on the LC in male and female mice, focusing on sensory thresholds, emotional states, and cognitive function. Nerve injury caused immediate sensory hypersensitivity in both sexes, while depressive-like behaviours and cognitive impairments emerged only after prolonged injury. Interestingly, anxiety-like behaviours and heightened fear conditioning were exclusive to males, which correlated with specific structural and functional changes in the male LC, such as increased noradrenergic cell counts, larger somato-dendritic volume, and greater neuron excitability. In females, however, neuronal excitability was reduced. Chemogenetic inhibition of LC neurons alleviated depressive- and anxiety-like behaviours, as well as fear conditioning, but only in males.

Overall, neuropathic pain induces distinct behavioural and neurobiological responses in males and females, challenging traditional views on female vulnerability to pain-induced anxiety and highlighting the need for sex-specific LC-targeted therapies.

## Introduction

Chronic pain affects 20-30% of adults globally, with over half also suffering from depression and anxiety (Lerman et al., 2015). Women experience higher rates of chronic pain and stress-related disorders, often alongside depression and anxiety (Gureje et al., 2008). Despite this, most preclinical studies on chronic pain comorbidities have focused on male rodents, thereby reflecting primarily male biological mechanisms. These findings underscore the need to include both male and female rodent models in chronic pain research to better understand sex differences and improve the relevance of findings.

Some studies have reported sexual dimorphism in several brain structures, including the Locus Coeruleus (LC) in both humans and rodents (Joshi and Chandler, 2020). The LC, a noradrenergic (NE) nucleus located in the brainstem, integrates signals related to attention, anxiety, stress response, arousal/sleep, learning and memory, sensory processing, pain, and reward processing (Chandler et al., 2019). In naïve rodents, the LC in females receives more stress-related afferents from regions such as the central amygdala and the bed nucleus of the stria terminalis (Valentino and Bangasser, 2016). Furthermore, female rats exhibit a more complex dendritic morphology in the LC compared to males (Bangasser et al., 2011; Luque et al., 1992). Accordingly, our previous studies have identified pronounced sex differences in the LC structure and LC-related behaviours in naïve mice (Mariscal et al., 2023). In male rodents experiencing chronic neuropathic pain, caused by disease or damage to the somatosensory system, evidence suggests that, as time passess after injury, increased LC activity leads to heightened anxiety, aversive learning and memory, and behavioural despair, all of which closely mirror symptoms of psychiatric stress-related disorders (Camarena-Delgado et al., 2022; Llorca-Torralba et al., 2022). Consequently, chronic pain evolves from a purely sensory experience into a complex emotional and cognitive burden, deeply intertwined with mental health that can be influenced by sex. Therefore, we examine sex differences in the LC of nerve-injured animals and explore their implications for stress-related behaviours.

## Results

### Behavioural sex differences in nerve-injured mice

As expected, nerve injury significantly increased mechanical and thermal hypersensitivity in CCI-2/3w and CCI-7/11w male and female mice (Figure 1B; Figure 1–figure supplement 1A, B). Depressive-like behaviour assessed via the TST revealed that while CCI-2/3w mice did not show behavioural despair, both male and female CCI-7/11w mice exhibited increased immobility (Figure 1C) and clasping behaviours (Figure 1–figure supplement 1C, D) when compared to their respective sham controls. Consistent findings were observed in the FST in CCI-7/11w mice (Figure 1–figure supplement 1E).

**Figure 1.**
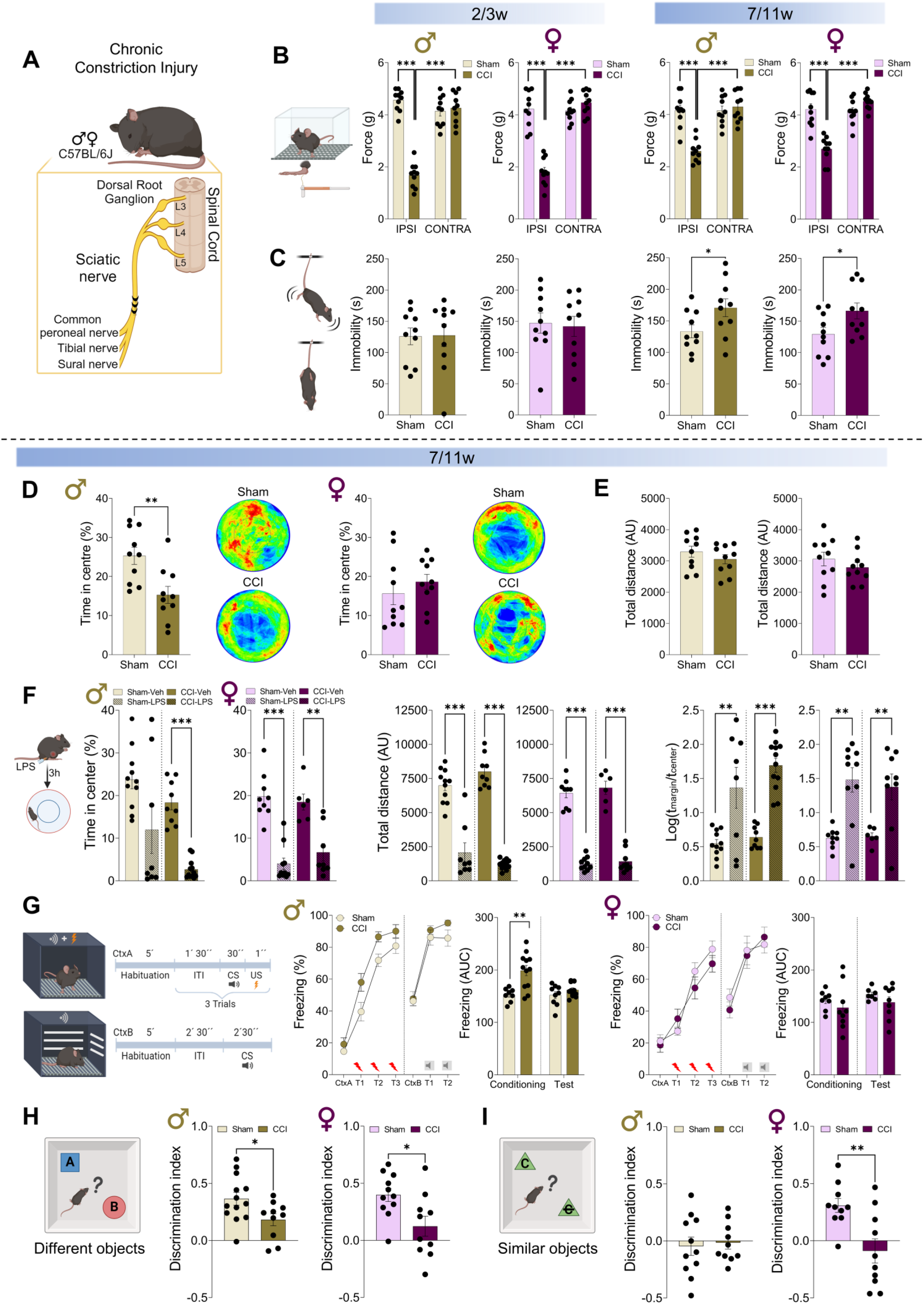
Sensory, emotional and cognitive consequences of neuropathic pain in male and female C57BL/6J mice. **(A)** Schematic representation of the chronic constriction injury (CCI) surgery for subsequent behavioural experiments in male and female C57BL/6J mice. **(B)** Mechanical hypersensitivity measured using von Frey filament stimulation (0 to 5 g, in 10 s) in male and female mice, tested 2-3 weeks or 7-11 weeks post-surgery. **(C)** Depressive-like behaviour assessed by the tail suspension test (TST), represented by immobility time (in seconds) for both male and female mice over the testing period. **(D)** Impact of neuropathic pain (CCI-7/11w) on anxiety, evaluated using the open field (OF) test and represented as the percentage of time spent in the central zone in both male and female mice, along with representative heatmaps of activity and **(E)** locomotor activity quantified as the total distance travelled (in Arbitrary Units, AU). **(F)** Stress-induced anxiety via acute lipopolysaccharide (LPS) systemic administration in the OF test, represented as the percentage of time spent in the central zone, total distance travelled and thigmotaxis, expressed as Log (time in margin / time in centre) for male and female mice. **(G)** Response to an aversive stimulus in the fear conditioning (FC) test. Schematic representation of the FC procedure for the different phases: day 1 (Context A, Conditioning) and day 2 (Context B, Test). The percentage of freezing represented during conditioned stimulus (CS) across trials and the area under the curve (AUC) of the percentage of freezing over time. **(H)** Schematic representations of the novel object recognition (NOR) experimental design to assess short-term memory (STM, learning index). Graph representing the discrimination index (DI) between objects following the STM protocols using a novel object (represented as B) that differed drastically from the familiar object (represented as A), and **(I)** using a novel object (represented as C crossed out) which presented strong similarity with the familiar one (represented as C) in both male and female mice. Data are presented as mean ± SEM of *n* = 7-13 mice per group: \*\*\**P* < 0.001 versus contralateral paw and sham counterparts, two-way ANOVA followed by Tukey’s post hoc test; \**P* < 0.05, \*\**P* < 0.01, \*\*\**P* < 0.001 versus sham or vehicle treatment, student’s t-test. IPSI, ipsilateral paw; CONTRA, contralateral paw; ITI, inter-trial interval; US, unconditioned stimulus.

Emotional impairments related to neuropathic pain were detected 7-11 weeks after nerve injury, prompting further assessment of emotional and cognitive behaviours at this time point. Anxiety-like behaviour, evaluated using the OF test, revealed that CCI-7/11w male mice spent significantly less time in the centre of the arena compared to sham males. In contrast, no differences were observed between sham and CCI-7/11w female mice (Figure 1D). Locomotor activity was similar across all groups, excluding the possibility of motor impairments (Figure 1E). To assess the impact of LPS administration, LPS-treated animals were tested in the OF test. These animals spent a reduced percentage of time in the centre zone and displayed decreased locomotor activity. The thigmotaxis ratio further confirmed an increased preference for the edges of the arena in LPS-treated animals, irrespective of sex or surgery (Figure 1F; Figure 1–figure supplement 2A, B), indicating heightened anxiety-like behaviour. In the fear conditioning test, CCI-7/11w male mice developed a higher percentage of freezing during the conditioning phase compared to sham males, while no differences were observed in CCI-7/11w female mice. In the testing phase, neuropathic and sham animals reached the same level of conditioning (Figure 1G).

In the classical NOR task, both CCI-7/11w male and female mice exhibited impaired recognition of the novel object, indicated by a decreased DI (Figure 1H; Figure 1–figure supplement 2C-G). In the NOR test assessing the ability to discriminate between two similar objects, both sham and CCI-7/11w males displayed a lower DI, reflecting a reduced ability to detect subtle differences. While sham females were able to recognize the novel object, this ability was significantly diminished in CCI-7/11w female mice, evidenced by a decreased DI (Figure 1I; Figure 1–figure supplement 2H-L).

### Structural, molecular and electrophysiological sex differences in the LC

Immunohistochemistry (Figure 2A; Figure 2–figure supplement 1A) revealed a significant increase in the number of TH+ cells in the LC of CCI-7/11w male mice, without changes in CCI-2/3w males. Additionally, no changes were observed in any female group (Figure 2B). Volumetric studies (Figure 2C; Figure 2–figure supplement 1B, C) revealed similar noradrenergic somatic volume between groups along neuropathy (Figure 2D). Furthermore, CCI males exhibited a significantly greater somatodendritic volume, attributed to an increase in dendritic volume, which persisted in both CCI-2/3w and CCI-7/11w groups. In contrast, nerve injury did not induce any changes in the LC of CCI females at any time point (Figure 2E, F).

**Figure 2.**
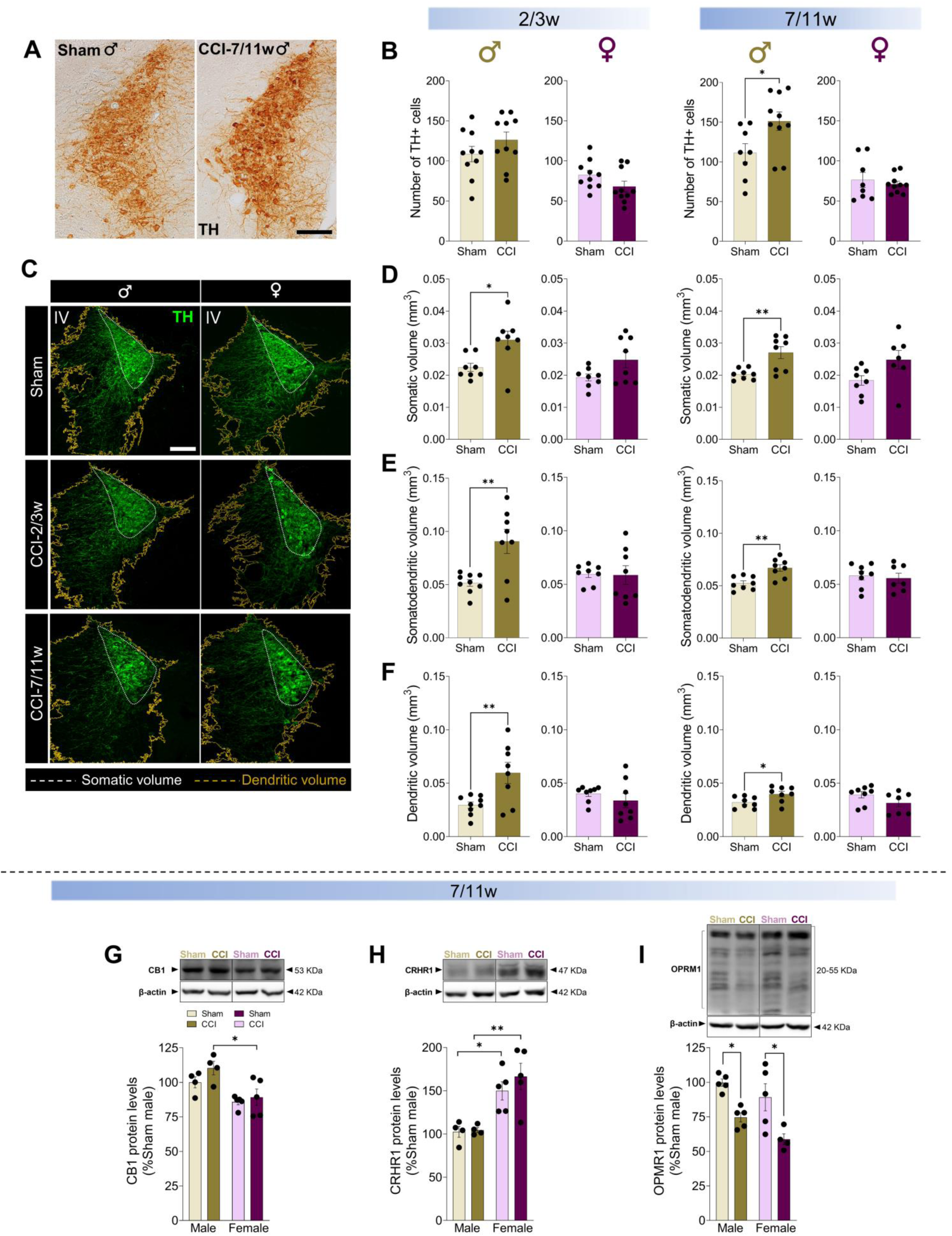
Quantification of TH+ cells and LC volume in male and female C57BL/6J mice after neuropathy. **(A)** Representative images of the LC of sham and CCI-7/11w male mice stained for TH using DAB. **(B)** Quantification of TH+ cell number in the LC of male and female C57BL/6J mice at 2-3 weeks and at 7-11 weeks post-surgery. **(C)** Representative confocal images illustrating the area occupied by the soma (white line) and dendrites (yellow line) used for volume estimation through TH immunofluorescence in the LC. **(D)** Volume occupied by the soma, **(E)** somatodendrites and **(F)** dendrites of the entire LC at 2-3 weeks and 7-11 weeks post-surgery in male and female mice. **(G)** Western blot analysis of cannabinoid receptor 1 (CB1), **(H)** corticotropin-releasing hormone receptor 1 (CRHR1) and **(I)** mu-opioid receptor 1 (OPMR1) protein levels in the LC of male and female mice subjected to CCI-7/11w. Data are presented as mean ± SEM of *n* = 10 LC per group for DAB staining, 7-10 LC per group for immunofluorescence, and a pool of LC from 5 mice per group for western blot analysis: \**P* < 0.05, \*\**P* < 0.01 versus sham, student’s t-test; \**P* < 0.05, \*\**P* < 0.01, two-way ANOVA followed by Tukey’s post hoc test. Scale bar = 100 μm. IV, fourth ventricle; CCI, chronic constriction injury; TH, Tyrosine Hydroxylase; DAB, 3,3′-diaminobenzidine tetrahydrochloride.

Molecular assessments revealed no significant changes in CB1 and CRF1 receptors levels in the LC associated with neuropathic pain in either males or females (Figure 2G, H). However, neuropathy was found to reduce OPMR1 protein levels in the LC of both CCI-7/11w male and female mice (Figure 2I). Furthermore, statistical analysis of sex differences revealed a reduction in CB1 receptor levels and an increase in CRHR1 receptor levels in the LC of females (Figure 2G, H).

Whole-cell patch clamp recordings of LC neurons were conducted in sham and CCI-7w mice. Results showed similar membrane resting potential, membrane capacitance and resistance in both sexes (Figure 3A, B). Resting IRK currents were similar in sham and CCI-7w males but significantly higher in CCI-7w females compared to sham females (Figure 3C). Further assessment of hyperpolarization and firing frequency in response to negative and positive current injections (25 pA steps) under current clamp mode showed that CCI-7w males exhibited a higher firing frequency than sham males at the same injected current, and consequently, lower rheobase current (Figure 3D). In contrast, CCI-7w females demonstrated lower excitability, and a higher rheobase compared to sham females (Figure 3F). Consistent with the IRK results, hyperpolarizing steps induced similar potentials in males (Figure 3E). However, CCI-7w females exhibited less pronounced hyperpolarized membrane deflections compared to sham females (Figure 3G).

**Figure 3.**
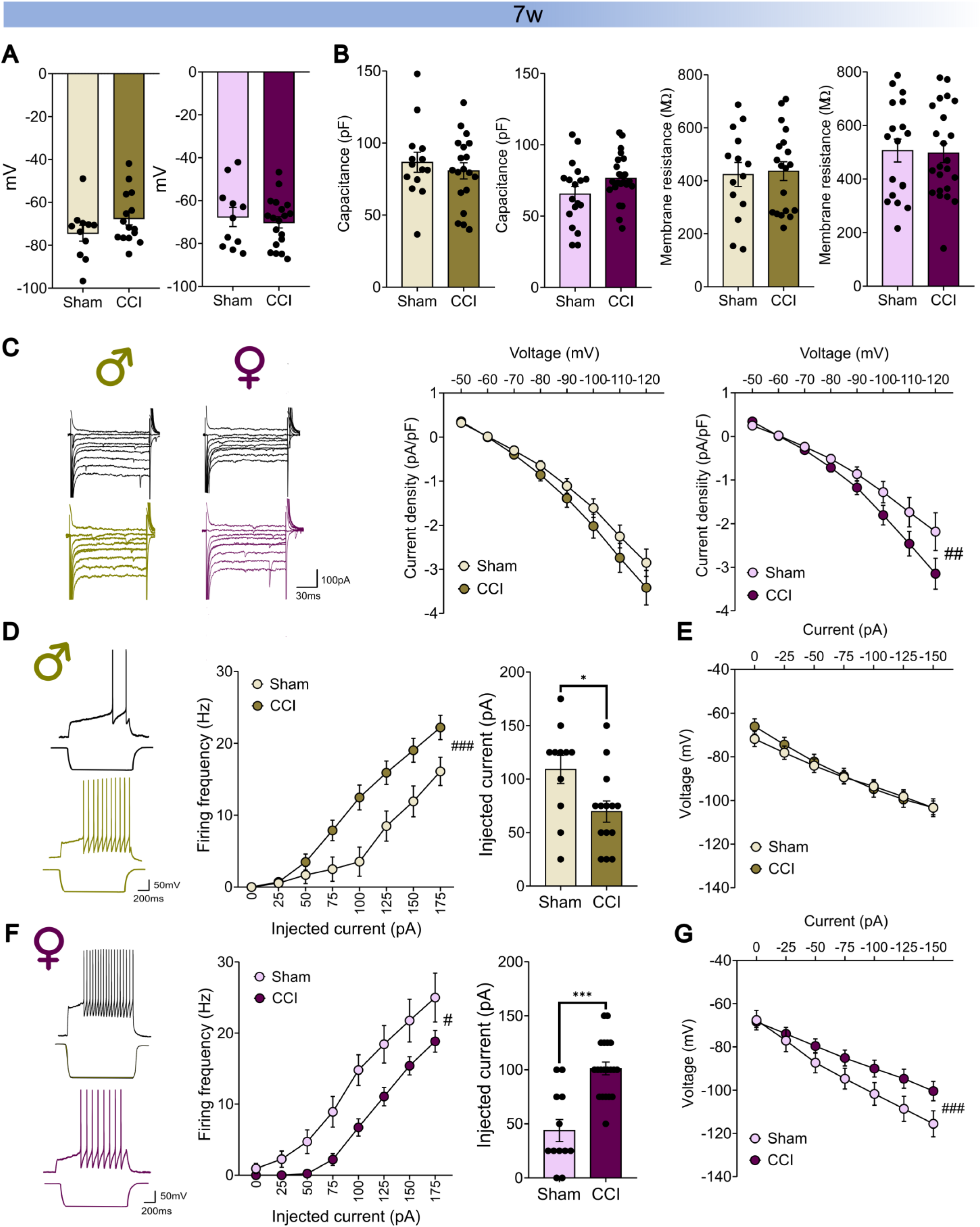
Evaluation of the passive properties and excitability of LC neurons in male (green) and female (purple) neuropathic mice. Graphs showing membrane resting potentials **(A)**, membrane capacitance and resistance **(B)**. **(C)** Representative examples and graphs of the IRK currents. **(D)** Representative examples of the voltage responses of identified LC neurons to current injection of + 100 and − 100 pA, respectively in sham and CCI-7w male mice. Graphs showing the activity driven in response to the injection of positive currents (+ 25 pA steps) and rheobase. **(E)** Graph showing the voltage deflections in response to the injection of negative currents (− 25 pA steps) in sham and CCI-7w male mice. **(F)** Representative examples of the voltage responses of identified LC neurons to current injection of + 100 and − 100 pA, respectively in sham and CCI-7w female mice. Graphs showing the activity driven in response to the injection of positive currents (+ 25 pA steps) and rheobase. **(G)** Graph showing the voltage deflections in response to the injection of negative currents (− 25 pA steps) in sham and CCI-7w female mice. Data are presented as the mean ± SEM of *n* = 5-6 mice per group: \**P* < 0.05, \*\*\**P* < 0.001 vs male, repeated-measures ANOVA (surgery*current factor): #*P* < 0.05, ##*P* < 0.01, ###*P* < 0.001. CCI, Chronic constriction injury; IRK, Inwardly rectifying potassium.

### Sex differences in the chemogenetic inhibition of LC neurons in nerve-injured mice

The sex-specific effects of LC noradrenergic neuron inhibition on emotional behaviours were assessed (Figure 4A, B; Figure 4–figure supplement 1A). In the TST, chemogenetic inhibition reduced immobility and increased latency to immobility in CCI-7/11w males, while no differences were observed in CCI-7/11w females (Figure 4C; Figure 4–figure supplement 1B). Anxiety-like behaviour, assessed using the OF test, revealed that chemogenetic inhibition of the LC increased the time spent in the central zone in CCI-7/11w males, an effect not observed in females. Furthermore, locomotor activity was similar across all groups (Figure 4D). In the fear conditioning test, chemogenetic inhibition of LC neurons resulted in a reduction of the percentage of freezing during the conditioning phase in CCI-7/11w males, while no differences were observed in CCI-7/11w females. No changes were seen in the testing phase (Figure 4E).

**Figure 4.**
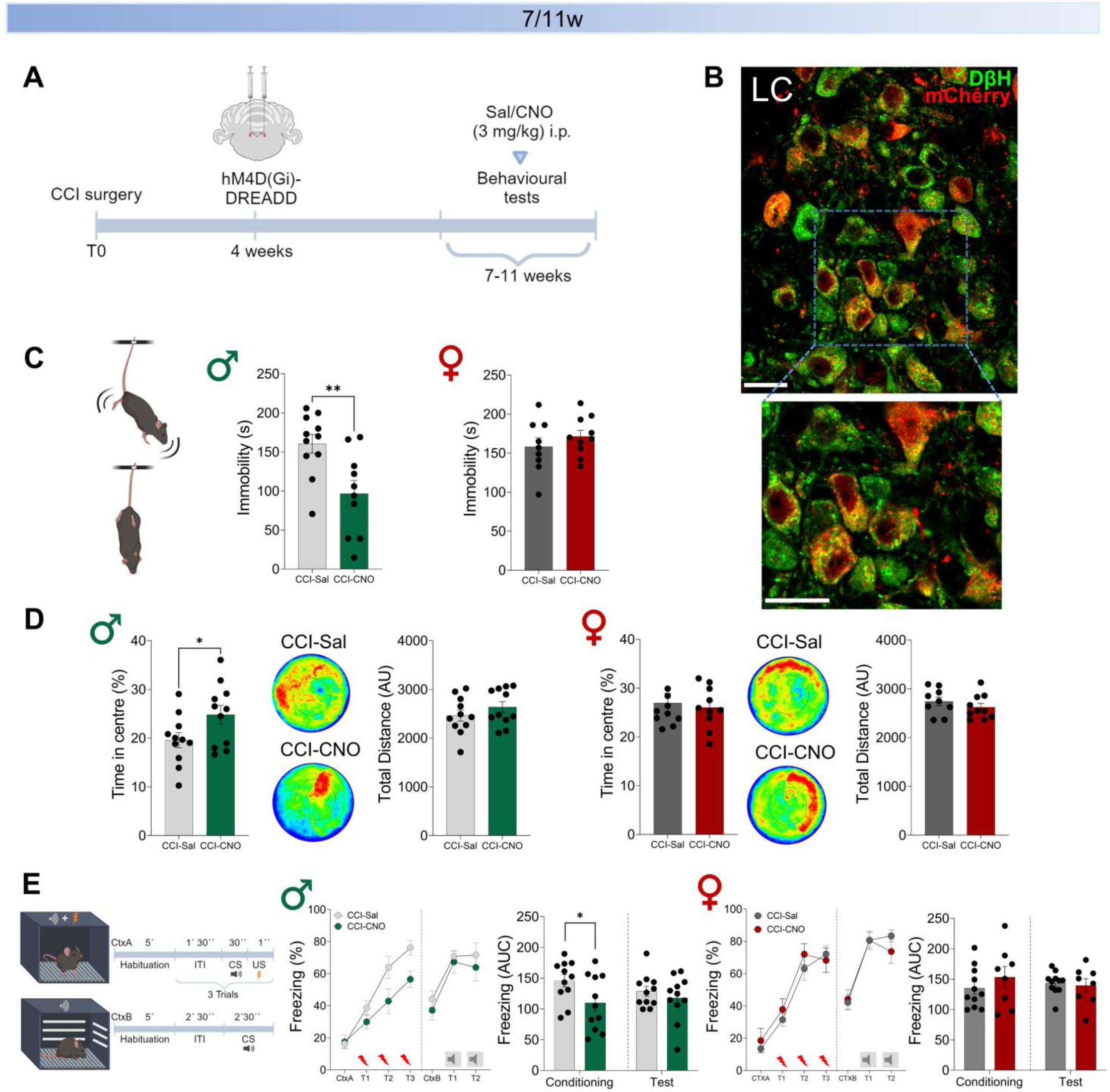
Effect of chemogenetic bilateral inhibition of noradrenergic neurons in the LC on depressive-like behaviour, anxiety and fear after 7-11 weeks of nerve injury in male and female TH:Cre mice. **(A)** Design of the experimental timeline. **(B)** Representative immunofluorescence image showing mCherry expression in noradrenergic LC neurons (red = mCherry, green = DβH) in TH:Cre mice (scale bar = 20 µm). **(C)** Depressive-like behaviour evaluated by the tail suspension test (TST), represented by immobility time (in seconds). **(D)** Anxiety-like behaviour represented as the percentage of time spent in the central zone in the open field (OF) test, along with representative heatmaps of activity, and the total distance travelled during the test (in Arbitrary Units, AU). **(E)** Response to an aversive stimulus in the fear conditioning (FC) test. Schematic representation of the FC procedure for the different phases: day 1 (Context A, Conditioning) and day 2 (Context B, Test). The percentage of freezing represented during the conditioned stimulus (CS) across trials, and the area under the curve (AUC) of the percentage of freezing over time. CNO (3 mg/kg) or saline was administered intraperitoneally 30 minutes prior to each behavioural test. Data are presented as the mean ± SEM of *n* = 8–11 mice per group: \**P* < 0.05, \*\*\**P* < 0.001 vs CCI-CNO, student’s t-test. CCI, Chronic constriction injury; CNO, Clozapine-N-oxide; ITI, inter-trial interval; US, unconditioned stimulus; DβH, dopamine β-hydroxylase.

## Discussion

This study examines the effects of nerve injury on the LC and related behaviours in mice, highlighting significant differences between sexes. Nerve injury reduced pain thresholds in both males and females at 2-3 and 7-11 weeks post-injury (Llorca-Torralba et al., 2022; Yalcin et al., 2011). While male mice exhibited both depressive- and anxiety-like behaviours following long-term neuropathic pain, female mice displayed depressive-like behaviour without a corresponding increase in anxiety. This aligns with findings from our systematic review and meta-analysis, which revealed a high prevalence of depression- and anxiety-like behaviours in neuropathic animals compared to sham controls, particularly in the FST and TST. However, a subgroup analysis noted substantial variation in the impact of neuropathic pain, influenced by factors such as species, strain, and model type, suggesting that specific conditions may promote the development of depression and/or anxiety-like behaviours (de la Rosa et al., 2024). A notable sex disparity emerged in our study, with females showing lower anxiety-like responses compared with males. Although this finding is somewhat counterintuitive given the higher prevalence of anxiety disorders and chronic pain in women (Tsang et al., 2008), it is supported by our experimental data, which showed that neuropathic males exhibited a higher percentage of freezing behaviour during the conditioning phase, consistent with previous studies in male neuropathic rats (Llorca-Torralba et al., 2019b). In contrast, anxiety-like behaviours were not heightened in neuropathic females. One potential explanation is that females might have reached the maximum anxiogenic response detectable in the OF test, but this seems unlikely since LPS treatment increased anxiety in both sexes. Alternatively, female neuropathic mice might require different stimuli to enhance anxiety or might employ alternative coping mechanisms involving distinct neural circuits for processing pain-related anxiety. Regarding non-emotional learning and memory, neuropathy led to deficits in the NOR test for both sexes, consistent with previous data. Recently, we reported that naïve female mice initially outperformed males in tasks requiring differentiation of objects with subtle differences (Mariscal et al., 2023). However, this cognitive advantage was lost following the induction of CCI, indicating that nerve injury not only causes general cognitive deficits but also specifically impairs the advanced cognitive abilities seen in naïve females.

Behavioural differences between sexes may be partly attributed to sex-specific structural and functional adaptations in the LC. Previous research has shown that naïve female mice have fewer TH+ cells but greater dendritic volume in the LC, along with higher CRHR1 expression and lower CB1 protein levels (Bangasser et al., 2016; Mariscal et al., 2023; Wyrofsky et al., 2018). These factors contribute to increased LC excitability, suggesting greater connectivity to emotional-related brain areas, which may underlie the heightened emotional arousal and anxiety observed in naïve female mice (Mariscal et al., 2023). Under neuropathic conditions, although both sexes displayed decreased protein expression of LC mu-opioid receptors, other adaptations were notably sex-specific. Nerve-injured males showed a significant increase in TH+ cell numbers and somatodendritic and dendritic volume, along with heightened excitability of LC neurons, consistent with previous findings (Llorca-Torralba et al., 2019a). In contrast, females exhibited no significant changes in LC structure and reduced neuronal excitability in patch clamp studies, suggesting that neuropathy sensitises LC neurons in males while desensitises them in females. Chemogenetic experiments further supported these sex differences. DREADD-mediated inhibition of LC neurons reversed neuropathic pain-induced negative consequences in males but had no effect on female behaviour, aligning with previous studies showing that increased LC activity impaired cognitive performance in males but not in females (Cardenas et al., 2021). These findings highlight the distinct neural mechanisms underlying emotional and cognitive responses to neuropathic pain between sexes, challenging conventional theories of female biological vulnerability to chronic pain and stress-related disorders.

Our study highlights the need for research into sex-specific pain pathways, especially in females, where mechanisms differ from those in males. Nerve injury sensitises the noradrenergic LC in males, increasing anxiety-like behaviour and aversive learning, a response not seen in females. Optogenetic and chemogenetic activation of the LC or its projections to the basolateral amygdala exacerbates anxiety and depression-like behaviours in sham/naïve male animals (Llorca-Torralba et al., 2022; Llorca-Torralba et al., 2019b; McCall et al., 2017), suggesting that females might be more vulnerable to the adverse effects of LC activation. This aligns with clinical findings where women show better responses to selective serotonin reuptake inhibitors (SSRIs) and lower tolerance to tricyclic antidepressants compared to men (LeGates et al., 2019). Our results emphasise the importance of adopting sex-specific approaches when targeting LC activity in chronic pain, recognising distinct emotional and cognitive outcomes based on sex.

## Materials and methods

### Animals and experimental design

Experiments were conducted on adult (8 – 12 weeks of age) male and female wild-type or tyrosine hydroxylase-Cre (TH:Cre) transgenic C57BL/6J mice (founders provided by EMMA INFRAFRONTIER, EM:00254) (INFRAFRONTIER Consortium, 2015). Mice were housed in separate rooms according to their sex under standard laboratory conditions (22°C, 12-h light-dark cycle) with *ad libitum* access to food and water. All procedures were approved by the Committee for Animal Experimentation of the University of Cádiz and the UPV/EHU (M20/2021/234 and M30/2021/235), in compliance with the European Commission’s Directive (2010/63/EU), Spanish Law (RD 53/2013) and ARRIVE guidelines.

Experimental time points set at 2-3 weeks (CCI-2/3w) or 7-11 weeks (CCI-7/11w) post-nerve injury. Mice were randomly assigned to experimental cohorts, and no additional strategies were employed to minimize potential confounders. Cohort #1 and #2: sham and CCI mice (n = 10 per group) underwent von Frey and TST at 2-3w and 7-11 weeks post-surgery, respectively. Cohort #3 and #4: sham and CCI mice (n = 10 per group) underwent acetone test and FST at 2-3w and 7-11 weeks post-surgery, and tissue for immunohistochemistry, immunofluorescence and western blot analysis was collected at least 1 week after the stressful event. Cohort #5: sham and CCI mice (n = 10) were subjected to the OF test and the plantar test 7-11 weeks post-surgery. Cohort #6: sham and CCI mice (n = 7-12 per group) were used to evaluate stress-induced anxiety via LPS administration at 7-11 weeks post-surgery. Cohort #7: sham and CCI mice (n = 8-12 per group) were subjected to the FC test at 7-11 weeks post-surgery. Cohort #8: sham and CCI mice (n = 10-13 per group) performed both approaches in the NOR paradigm at 7-11 weeks post-surgery. Cohort #9: sham and CCI mice (n = 5-6 per group) were used for patch-clamp studies at 7 weeks post-surgery. Cohort #10: CCI-saline and CCI-CNO mice (n = 9-11 per group) were administered DREADDs and performed the OF test and the TST at 7-11 weeks post-surgery. Cohort #11: CCI-saline and CCI-CNO mice (n = 8-11 per group) performed the FC test at 7-11 weeks post-surgery. All animals were habituated to the testing room for 30 minutes prior experiments. All behavioural approaches were conducted under 13 lux lighting, and apparatuses were cleaned with 70% ethanol between animals, especially between sexes. Experiments and analyses were performed blind to the assigned group or treatment.

### Neuropathic pain model

Chronic constriction injury (CCI) of the sciatic nerve was used as a model for neuropathic pain (Figure 1A) (Bennett and Xie, 1988; Emery et al., 2011). Mice were anaesthetised on 4% isoflurane and maintained on 1.5-2.5% isoflurane in a mixture of O_2_ at a flow rate of 0.5 L/min. The left hind paw was shaved at the mid-thigh level, and after a skin incision, the gluteus superficialis and the biceps femoris muscles were separated using curved forceps. The left sciatic nerve was exposed proximal to the trifurcation, and three chromic gut (6-0) ligatures were loosely tied around the nerve at 1-mm intervals. The skin layers were subsequently closed using 5-0 silk thread. Sham-operated mice underwent the same procedure, except for nerve ligation.

### DREADD virus injection

Male and female TH:Cre CCI-7/11w mice were bilaterally injected into the LC with a cre-dependent DREADD inhibitor virus [AAV2/hSyn-DIO-hM4D(Gi)-mCherry, hM4D(Gi)-DREADD, from Virus Vector Core, Gene Therapy Center Vector Core at the University of North Carolina, USA (Llorca-Torralba et al., 2019b)]. Briefly, mice were anaesthetised and placed in a stereotaxic frame (Kopf Instruments). Bupivacaine (2 mg/kg) and lidocaine (3.5 mg/kg) were administered subcutaneously in the incision area prior to surgery. The DREADD virus (titer 3×10^12 vg/ml) was bilaterally injected at a volume of 0.5 µl into the LC (AP: −5.5 mm, ML: ±1 mm, DV: 3 mm) (Paxinos and Franklin, 2019), and the skin was closed using 5-0 silk thread. DREADD expression in the LC was verified at the end of the experiments in the LC and other noradrenergic areas as A5 and A7 (Figure 4–figure supplement 1A). Behavioural testing was carried out 3 weeks after virus injections to maximise vector expression, coinciding with the CCI-7/11w time point. Experimental groups included CCI-7/11w male and female mice receiving DREADD virus injections into the LC, were administered intraperitoneally with either saline or clozapine-N-oxide (CNO, 3 mg/kg) for chemogenetic LC inhibition, 30 minutes prior to each behavioural test with experimenters blind to the treatment.

### Sensory, emotional and cognitive assessment

#### Von Frey test

Mechanical hypersensitivity was assessed at CCI-2/3w and CCI-7/11w after injury using a Dynamic Plantar Aesthesiometer (Ugo Basile, Italy). After 30 minutes of habituation on a wire-mesh floor, a force ranging from 0 to 5 grams over 10 seconds was applied to the plantar surface of the hind paws. The withdrawal threshold (grams) was determined as the average of two measurements that elicited a rapid paw withdrawal (Emery et al., 2011).

#### Acetone test

The acetone evaporation test was used to assess cold hypersensitivity. Mice underwent a 30-minute habituation period in individual plastic cages placed on an elevated mesh floor. A 50 µl drop of acetone was applied to the plantar surface of each hind paw to induce evaporative cooling. The time spent lifting, licking, and/or shaking of the hindlimb was monitored for 60 seconds, and the time of response was calculated by averaging the results of three acetone applications (Catuneanu et al., 2019; Colburn et al., 2007).

#### Plantar test

Thermal hyperalgesia was evaluated by the Hargreaves’ method (Hargreaves et al., 1988). Mice were habituated for 45 minutes in individual plastic cages on an elevated glass surface, and a radiant heat source (Plantar test device, Ugo Basile, Italy) was applied to the plantar surface of each hind paw at a constant intensity, with a 30-seconds cut-off to prevent skin damage. Two measurements were made and the mean latency (in seconds) of the paw withdrawal was considered the thermal nociceptive threshold (Alba-Delgado et al., 2012).

#### Tail suspension test (TST)

Mice were suspended by the distal end of their tails and raised 20 cm above the floor (Berrocoso et al., 2013). The latency to the first episode of immobility and the total duration of immobility, defined as the state of hanging by the tail without any active movements, were analysed during the 6-minute test session at CCI-2/3w and CCI-7/11w after injury. Additional behaviours were also assessed: a) climbing - the animal climbs on its tail; b) swinging - the animal moves its body from side to side c) curling - the animal performs twisting movements with its body actively; d) clasping - the animal retracts its hind limbs towards the abdomen.

#### Forced swimming test (FST)

The FST was used to assess depressive-like behavior (Berrocoso et al., 2006; Porsolt et al., 1977). Each animal was placed in a glass cylinder (10 cm diameter x 18 cm height) filled with water to a depth of 10 cm, at a temperature of 22 ± 1°C, for a 6-minute test. The sessions were recorded and the latency to the first bout of immobility and the time spent immobile during the last 4 minutes of the testing period were analysed. Immobility behaviour was defined as the minimal movements necessary to keep the animal’s head above water.

#### Open field (OF) test

Mice were individually placed in the centre of a 40-cm circular enclosure, and allowed to move freely for 10 min while being recorded. The centre of the arena was defined as an area comprising 50% of the total arena. The percentage of time spent in the centre (as an indicator of anxiety-like behaviour) and the total distance travelled (indicating locomotor activity) were analysed using the SMART video-tracking software (Panlab, S.L., Barcelona, Spain) at CCI-7/11w after injury. Thigmotaxis was calculated as the Log (time in margin / time in centre) (Wurzman et al., 2015).

#### Stress-induced anxiety via Lipopolysaccharide (LPS) administration

LPS (*Escherichia Coli* 0111:B4, Sigma-Aldrich) in sterile PBS was administered intraperitoneally at a single dose of 100 µg/kg (Mulvey et al., 2018). LPS-induced anxiety was assessed 3 hours post-injection using the OF test at CCI-7/11w, as previously described.

#### Fear conditioning (FC) test

In the fear conditioning test, a tone (30 seconds, 5 kHz, 50 dB) was used as a conditioned stimulus (CS), and a foot shock (1 second, 0.1 mA) was used as an unconditioned stimulus (US). Freezing was evaluated during the conditioned stimuli, as the absence of all movement except that necessary for respiration (Curzon et al., 2009). CCI-7/11w animals were fear conditioned (day 1) and context tested (day 2) in a Shuttle Box using two context options: Context A (CtxA) involves a black chamber with white light and scented with a 75% ethanol solution; Context B (CtxB) involves a black/white chamber with red light and scented with soapy water. Prior to conditioning and context tested, mice were habituated to the apparatus for 5 minutes. On the day of conditioning, mice were given 3 CS-US pairings during the CtxA. The inter-trial interval (ITI) was 90 seconds. The conditioned mice were tested for fear responses (freezing) 24 hours after conditioning. For this test, mice were assessed in CtxB and exposed to 1 CS of 150 seconds. The percentage of freezing was calculated manually by using SMART software and represented during each CS.

#### Novel Novel Object Recognition (NOR) test

The NOR paradigm was employed to assess short-term memory (STM) as a learning index, using two distinct approaches at CCI-7/11w. Firstly, a classical NOR protocol with a new object completely different in shape, colour and texture from the familiar one (Figure 1–figure supplement 2C). The second approach aimed to evaluate the ability to discern subtle differences between objects (Figure 1–figure supplement 2H) (Mariscal et al., 2023). The test was conducted in a 45 cm x 45 cm squared enclosure. Mice underwent a 10-minute habituation phase in the absence of objects, followed by a 10-minute training phase where they were allowed to explore two identical objects placed in opposite corners of the arena. To evaluate short-term memory, mice were returned to their home cages for a 1-hour delay period. Subsequently, they were reintroduced to the arena for a 10-minute test phase, with one of the familiar objects replaced by a novel object. The analysis measured preference for identical objects (%), total number of interactions with objects, latency to the novel object, and the Discrimination Index (DI), calculated as (Tnovel - Tfamiliar) / (Tnovel + Tfamiliar), where T represents the time exploring each object (Suárez-Pereira and Carrión, 2015). Animals showing a greater preference for one of the two familiar objects during the training phase were excluded from the analysis.

### Immunohistochemistry and immunofluorescence

#### Perfusion and sample extraction

Mice were anaesthetised with 25% sodium pentobarbital and transcardially perfused with 0.9% saline solution followed by 4% paraformaldehyde (PFA) prepared in 0.1 M phosphate buffer saline (PBS). Brains were post-fixed in 4% PFA for 2 hours, then immersed in a 30% sucrose solution with 0.1% sodium azide in 0.1M phosphate buffer, and stored at 4°C for at least one overnight. Coronal sections (40µm) were cut using a freezing microtome, and preserved in cryoprotectant at 4°C until further analysis.

#### Structural and morphological studies

Free-floating immunohistochemistry was performed as previously described (Mariscal et al., 2023), using one of four series of 40µm-thick LC sections from 5 mice per group at CCI-2/3w and CCI-7/11w. Firstly, endogenous peroxidase activity was blocked by incubating the sections with 0.3% H_2_O_2_ in PBS for 30 minutes, followed by a 1-hour incubation with 2.5% bovine serum albumin (BSA) in 0.1 M PBS containing 0.3% Triton X-100. Then, sections were incubated with primary anti-Tyrosine Hydroxylase antibodies (rabbit anti-TH, 1:1000; OPA1-04050, Millipore) for two nights at 4°C. After washing, the sections were incubated with biotinylated donkey anti-rabbit antibodies (1:200, Jackson ImmunoResearch Europe, UK) for 60 minutes at room temperature. Subsequently, the ultra-sensitive ABC peroxidase staining kit (1:1000, Thermo Scientific, Spain) and DAB (3,3’-diaminobenzidine tetrahydrochloride) were used to visualise immunostaining. Sections were mounted on glass slides, cleared in xylene, and coverslipped using DPX mounting medium. An Olympus BX60 microscope equipped with an Olympus DP74 camera was used to capture images with the same exposure and illumination settings for all groups. The number of noradrenergic cells in the LC was manually counted using the Cell Counter plugin in ImageJ (National Institutes of Health, Bethesda, Maryland).

Immunofluorescence analysis was conducted on all the 40µm-thick LC sections from a different set of 5 mice per group at CCI-2/3w and CCI-7/11w. The same primary antibody was used, followed by appropriate fluorophore-conjugated secondary antibodies (Donkey anti-rabbit Alexa 488, 1:1000; A-21206, Invitrogen). After mounting on glass slides with a hard-setting antifade medium (Dako, S3023), the fluorescent signal was observed and captured using a 20X oil immersion objective on a confocal microscope (Olympus FV1000) with consistent exposure settings. Imaging systematically covered the entire rostrocaudal extent of the LC (approximately - 5.32 mm to - 5.82 mm from Bregma). The LC volume was estimated based on Cavalieri’s principle (Michel and Cruz-orive, 1988) following the formula V = ΣA x T, where ΣA is the sum of the measured areas in all LC sections, and T is the distance between sections. Thus, the area occupied by the somas of the LC cells, as well as the somato-dendritic area, comprising the entire LC, were calculated in each 40 µm-spaced parallel LC section. These areas were automatically delineated using the wand tool in an 8-connected configuration mode to detect connected regions. Subsequently, the dendritic area was estimated by subtracting the soma area from the entire LC area. The representation of each area across all the LC sections was depicted and the area under the curve (AUC) was also calculated along the rostro-caudal axis. Results were expressed in mm^2^ for areas and mm^3^ for volumes.

#### DREADD expression in LC-noradrenergic neurons

To evaluate the selective expression of DREADD in the LC and its absence in the A5 and A7 noradrenergic nuclei (Figure 4–figure supplement 1A), cryosections were incubated with a primary antibody cocktail to detect red fluorescent proteins (rat anti-RFP (mCherry), 1:500, 5F8, ChromoTek) and dopamine β-hydroxylase (rabbit anti-DβH, 1:500, ab20948, Abcam), followed by appropriate fluorophore-conjugated secondary antibodies (Streptavidin Alexa Fluor 568 conjugate, 1:1000, S11226, Invitrogen; Donkey anti-rabbit Alexa 488, 1:1000, A-21206, Invitrogen). Images were captured and analysed for colocalization using a Zeiss LSM 900 Airyscan 2 confocal microscope. Mice with no DREADDs expression in the LC were excluded from the analysis.

### Western blot assay

The protein expression level of receptors that modulate the LC-noradrenergic system (Llorca-Torralba et al., 2019a; Wyrofsky et al., 2018), including cannabinoid receptor 1 (CB1), corticotropin-releasing hormone receptor 1 (CRHR1), and mu-opioid receptor 1 (OPRM1) were evaluated in the LC of CCI-7/11w mice by western blotting. LC samples from five mice per group were dissected and pooled for protein isolation. Tissue homogenization was carried out through sonication in 200 µl of PBS (pH=7) mixed with a protease inhibitor cocktail (Complete, Roche, 04693116001), followed by centrifugation at 12 000 rpm for 10 min at 4 °C. Supernatants were collected and quantified using the Bradford method, and then mixed with Laemmli sample buffer (BioRad, 1610737). 20 µg/µl were loaded and separated in 10% SDS-polyacrylamide gel electrophoresis (110 V). Subsequently, 0.2 µm nitrocellulose membranes (BioRad, 170459) were blocked in Tris-buffered saline supplemented with 0.1% Tween 20 and 5% BSA, and then incubated with the following antibodies overnight: CB1 (Invitrogen, PA1- 743, 1/250), OPMR1 (Invitrogen, 44-308G, 1/1000), and CRHR1 (Invitrogen, 720290, 1/1000). The next day, primary antibodies were washed, and membranes were incubated with their respective horseradish peroxidase-conjugated secondary antibodies for 1 h 30 min at room temperature and revealed using an ECL kit (Amersham Ibérica, 12644055). Blots were imaged using the ChemiDoc Imaging System (BioRad, 12003153) and quantified by densitometry with the NIH ImageJ software. Every densitometry is expressed as arbitrary units of optical density. β-actin (Sigma-Aldrich, A5441, 1/8000) was used as loading control (Kasai et al., 2011; MacDowell et al., 2017).

### Whole-Cell Patch-Clamp Recordings

Whole-cell patch-clamp recordings of LC neurons were performed at CCI-7w as previously described (Mariscal et al., 2023). Sham and CCI-7w mice (5-6 males and 6 females per group) were sacrificed by decapitation under deep anesthesia (4% isoflurane), brains were removed and transferred to ice-cold artificial cerebrospinal fluid (ACSF, pH 7.4) equilibrated with 95% O_2_ and 5% CO_2_, and containing (in mM): 250 sucrose, 26 NaHCO_3_, 1.25 NaH_2_PO_4_.H_2_O, 0.5 CaCl_2_.2H_2_O, 10 MgSO_4_.7H_2_O, 10 D-glucose. Coronal sections of the brain containing the LC (220 µm thick) were obtained with a vibratome (VT1200S; Leica Microsystems, Germany) and slices were incubated in warmed (30-35 °C) ACSF for at least 30 minutes before recording. Each slice was transferred to a recording chamber that was perfused continuously with oxygenated ACSF at 32–34 °C following our standard protocol (Miguelez et al., 2012; Rivera et al., 2021). LC neurons were visualized using infrared gradient contrast video microscopy (Eclipse workstation, Nikon) and with a 60X water-immersion objective (Fluor 60X/1.00 W, Nikon). The LC was identified as a dense and compact group of cells at the lateral border of the central gray and the fourth ventricle, just anterior to the genu of the facial nucleus. Recordings from individual LC neurons were obtained with pipettes (impedance, 3–6 MΩ) prepared from borosilicate glass capillaries (G150-4: Warner Instruments, Hamdem, CR, USA). The patch pipette was filled with a KGluconate-based solution containing (in mM): 130 KGluconate, 5 NaCl, 1 MgCl2.6H2O, 10 HEPES, 1 Na4EGTA, 2 MgATP, 0.5 NaGTP, and 10 Na2PCr. The junction potential between the electrode solution and the external media (empirically estimated as 13 mV) was not corrected, and electrode signals were low-pass filtered at 4 kHz and sampled at 20 kHz. LC neurons were identified by the presence of a resting inwardly-rectifying potassium (IRK) conductance by stepping the membrane potential from − 40 to − 120 mV in − 10 mV increments (100 ms/step) (Williams et al., 1988). In voltage clamp experiments, neurons were maintained at − 60 mV and the series resistance was monitored with steps of − 5 mV at the end of each recording. Data were discarded when the series resistance increased by > 20%. The average current response was analysed off-line and the cell capacitance (Cm) and membrane resistance (Rm) were calculated. In the current clamp mode, incremental currents from − 150 to + 300 pA were injected in 25 pA steps to explore the subthreshold and firing properties of the neurons. Off-line analysis was performed using pClamp V9.2 (Molecular Devices, San Jose, CA, USA).

### Statistical analysis

Data are presented as mean ± SEM. The sample size was estimated using the G*Power tool to detect a significant effect using two-tailed t-tests at an α level of 0.05, power of 0.80, and effect size based on the mean values from published reference studies on mechanical and thermal sensory thresholds in mice (Barthas et al., 2015; Li et al., 2022). Statistical analysis was performed using GraphPad Prism software (GraphPad Software 9.0.3, La Jolla, CA). Grubbs’ test was used to identify statistical outliers and the Shapiro-Wilk test to evaluate the normality of data distribution. For comparisons between two groups, unpaired Student’s t-test was used for parametric data, and Mann-Whitney U tests for non-parametric data. One-way, two-way, or repeated-measures ANOVA followed by Tukey’s post hoc tests were used for comparisons among multiple groups. Area under the curve (AUC) analysis was utilised for comparisons between groups across different time points. Chi-square test was employed for analysing frequency distribution. Statistical significance was set at P < 0.05.

## Data availability

All data generated during this study are included in the manuscript and supporting files. Source data and statistical analyses have been provided for all figures.

## Acknowledgements

The authors are sincerely grateful to Irene Suárez-Pereira, PhD., Jose Antonio Garcia Partida, M.Sc., and Elena Marín Álvarez, Higher Technician, from the University of Cádiz (Cádiz, Spain), for their excellent technical support. They also acknowledge the valuable assistance provided by the Central Services of Scientific and Technological Research, Health Sciences and Animal Research at the University of Cádiz. This research was conducted within the “Red Española de Investigación en Estrés/Spanish Network for Stress Research, RED2022-134191-T” financed by MICIU/AEI /10.13039/501100011033; Grant PRE2019-091106 and Grant PTA2021-019890-I funded by MICIU/AEI /10.13039/501100011033 and FSE+. Figures were created with BioRender.com.

---

**Figure 1–figure supplement 1.**
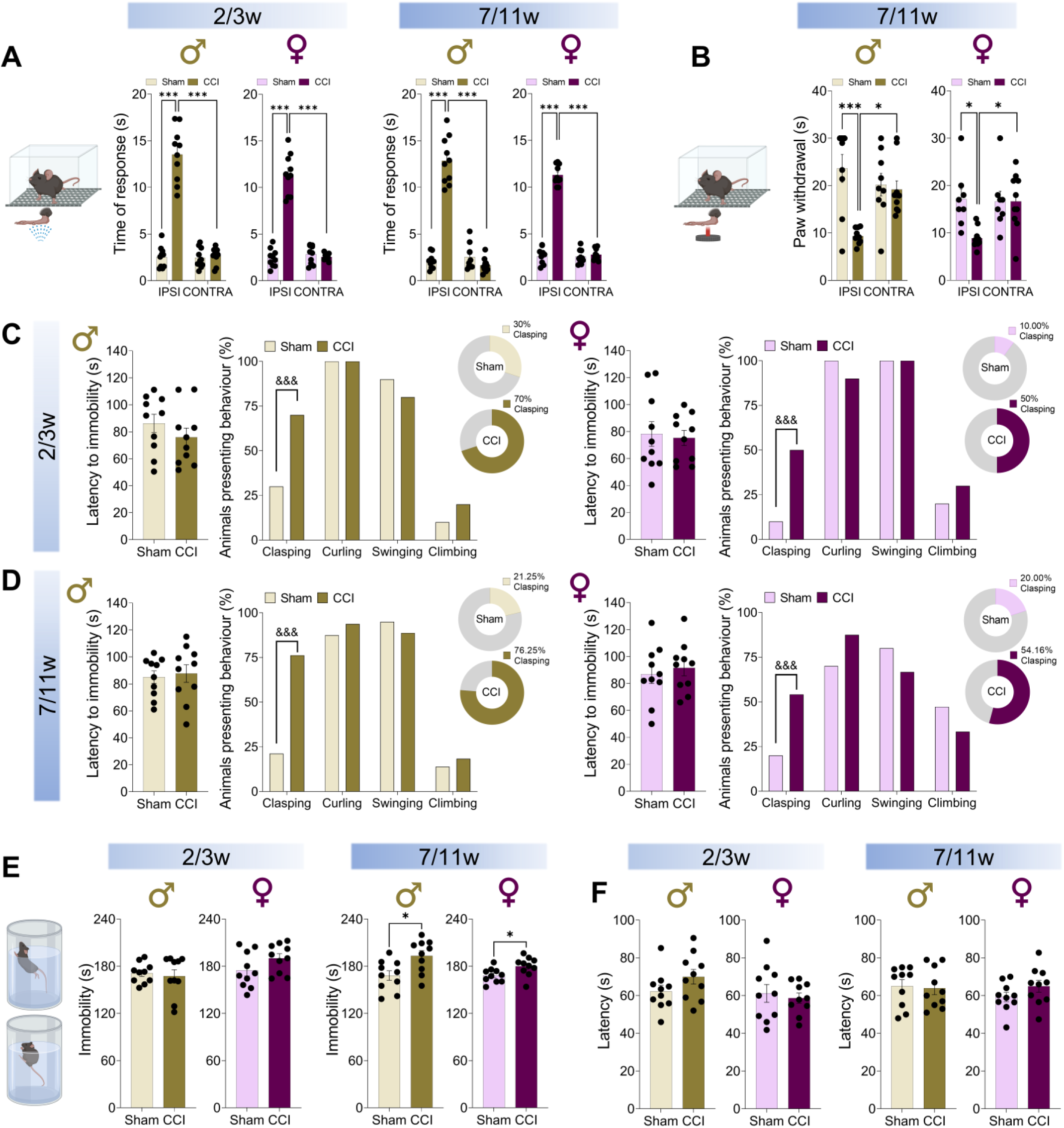
(A) Evaluation of cold hypersensitivity assessed using the acetone test, represented as the time of response (in seconds) in male and female mice at 2-3 weeks and 7-11 weeks post-surgery. **(B)** Thermal hyperalgesia evaluated in the plantar test, represented as the latency of paw withdrawal (in seconds) to a heat stimulus (cut-off 30 s) in CCI-7/11w male and female mice. **(C)** Graphs depicting the latency to immobility (in seconds) and the percentage of mice exhibiting clasping, curling, swinging and climbing behaviours in the TST at 2-3 weeks and **(D)** 7-11 weeks post-surgery. **(E)** Depressive-like behaviour expressed as immobility time in the FST and **(F)** the latency to immobility (in seconds) in male and female mice at 2-3 weeks and 7-11 weeks post-surgery. The data are presented as the mean ± SEM of *n* = 8-10 mice per group. \**P* < 0.05, \*\*\**P* < 0.001 versus contralateral paw or sham counterparts, two-way ANOVA followed by Tukey’s post hoc test; &&&*P* < 0.001 versus sham, Chi-squared test. \**P* < 0.05 versus sham, student’s t-test (See Supplementary Table 2 and 3). CCI, Chronic constriction injury; CONTRA, contralateral paw; FST, Forced swimming test; IPSI, ipsilateral paw; TST, Tail suspension test.

**Figure 1–figure supplement 2.**
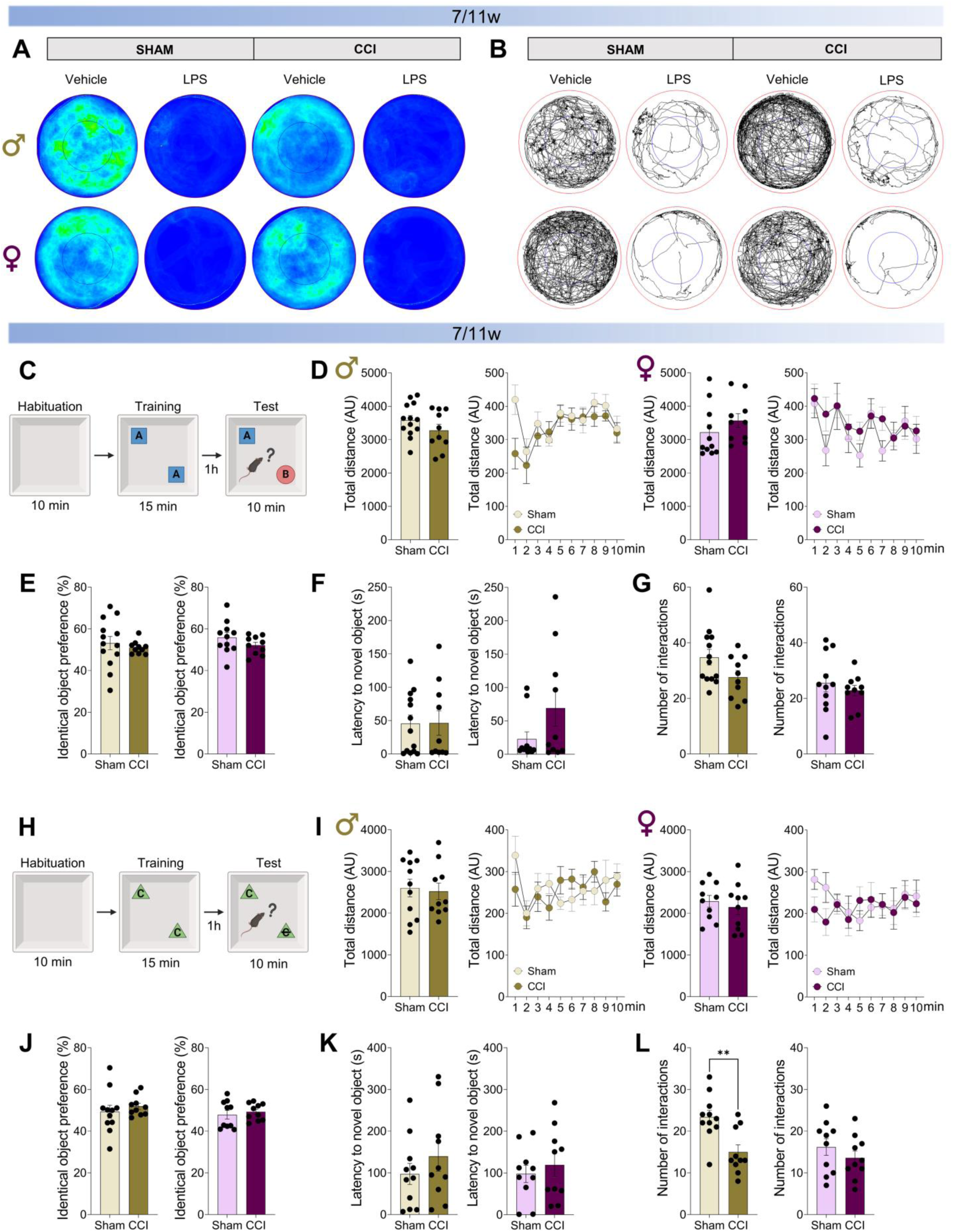
(A) Heatmaps of sham and CCI-7/11w male and female mice in the OF test 3 hour after vehicle or LPS intrapreritoneal administration, and **(B)** the respective trajectory traces. **(C)** Schematic representation of the novel object recognition (NOR) experimental design used to assess short-term memory (STM, learning index) using two completely different objects. **(D)** Graph showing the total distance travelled during the habituation phase of the test and the representation in 1-minute intervals (in arbitrary units, AU) for male and female mice at 7-11 weeks post-surgery. **(E)** Graph depicting the percentage of preference exploring identical objects (represented as A) during the 15-minute training phase of the test. **(F)** Latency to explore the novel object (represented as B) and the **(G)** number of interactions with objects during the test. **(H)** Schematic representation of the NOR experimental design used to assess STM using objects with minimal differences between them. **(I)** Graph showing the total distance travelled during the habituation phase of the test and the representation in 1-minute intervals (in arbitrary units, AU). **(J)** Graph depicting the percentage of preference exploring identical objects (represented as C) during the 15-minute training phase. **(K)** Latency to explore the novel object (represented as C crossed out) and the **(L)** number of interactions with objects during the test. The data are presented as mean ± SEM for *n* = 10-13 mice per group. \*\**P* < 0.01 versus sham, student’s t-test (See Supplementary Table 3). AU, Arbitrary Units; CCI, Chronic constriction injury; LPS, Lipopolysaccharide.

**Figure 2–figure supplement 1.**
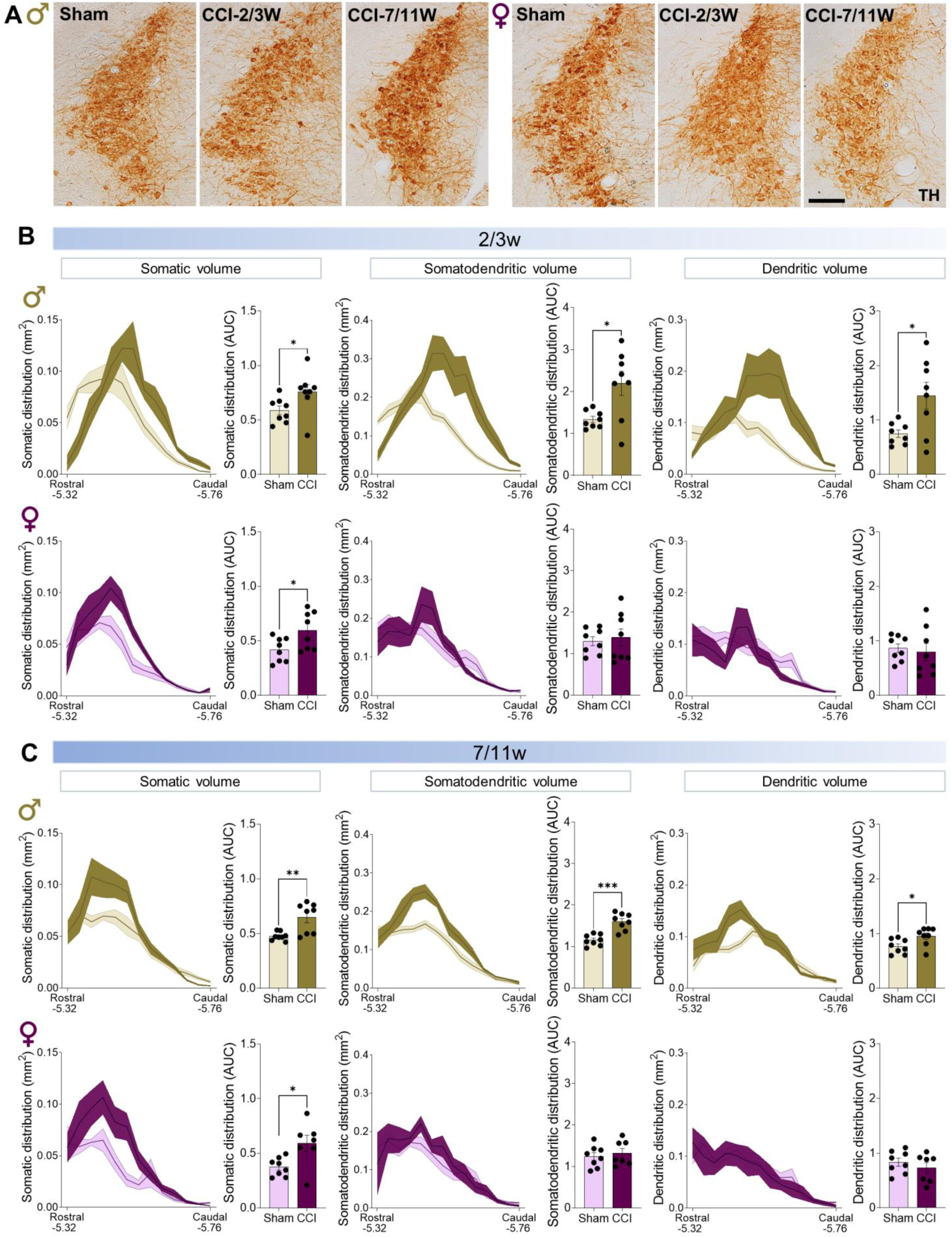
(A) Representative immunohistochemistry images of the LC stained for tyrosine hydroxylase (TH) by DAB in male and female mice. **(B)** Graphs depicting the distribution along the rostrocaudal axis of the area occupied by the soma, somatodendritic and dendrites in the entire LC (distances from Bregma in mm), and their representation as the area under the curve (AUC) in CCI-2/3w and in **(C)** CCI-7/11w male and female mice. The data are presented as mean ± SEM of *n* = 7-8 LC per group. \**P* < 0.05, \*\**P* < 0.01, \*\*\**P* < 0.001 versus sham, student’s t-test (See Supplementary Table 4). CCI, Chronic constriction injury; DAB, 3,3′-diaminobenzidine tetrahydrochloride. Scale bar = 100 µm.

**Figure 4–figure supplement 1.**
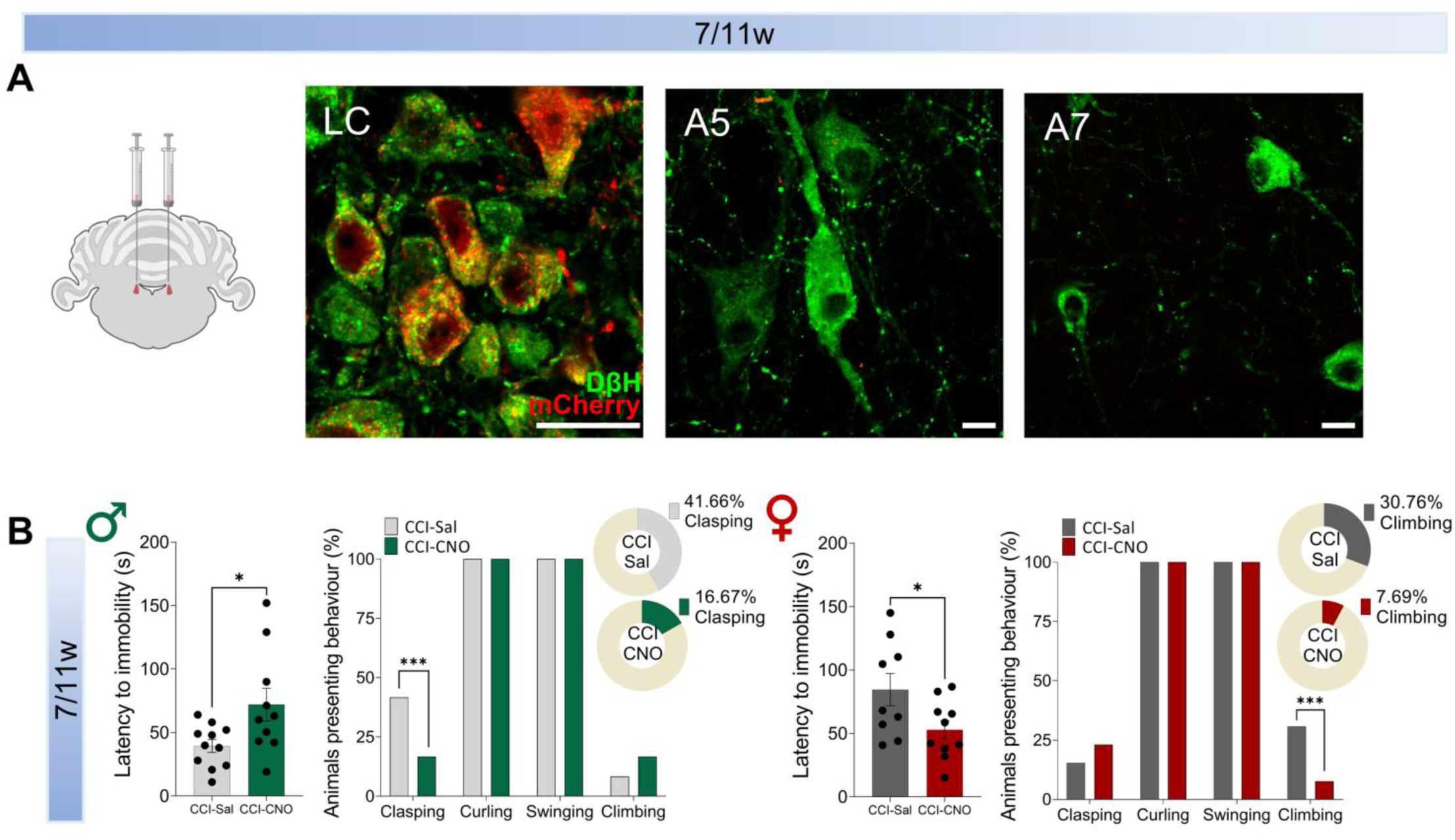
(A) Representative confocal images of immunofluorescence staining showing mCherry expression in noradrenergic LC neurons (red = mCherry, green = DβH) and other noradrenergic areas (A5 and A7) in TH:Cre male and female mice. **(B)** Latency to immobility (in seconds) and the percentage of clasping, curling, swinging and climbing behaviours in the TST in male and female mice at 7-11 weeks post-surgery. The data are presented as the mean ± SEM of *n* = 10-11 mice per group. \**P* < 0.05, student’s t-test; \*\*\**P* < 0.001, Chi-squared test (see Supplementary Table 6). CCI, Chronic constriction injury; CNO, Clozapine-N-oxide; DβH, dopamine β-hydroxylase; LC, Locus Coeruleus; Sal, Saline; TST, Tail suspension test. Scale bar = 20 µm.

## References

Alba-Delgado, C., Mico, J. A., Sánchez-Blázquez, P., Berrocoso, E., 2012. Analgesic antidepressants promote the responsiveness of locus coeruleus neurons to noxious stimulation: implications for neuropathic pain. Pain 153, 1438–1449.

Bangasser, D. A., Wiersielis, K. R., Khantsis, S., 2016. Sex differences in the locus coeruleus-norepinephrine system and its regulation by stress. Brain Res 1641, 177–188.

Bangasser, D. A., Zhang, X., Garachh, V., Hanhauser, E., Valentino, R. J., 2011. Sexual dimorphism in locus coeruleus dendritic morphology: a structural basis for sex differences in emotional arousal. Physiol Behav 103, 342–351.

Barthas, F., Sellmeijer, J., Hugel, S., Waltisperger, E., Barrot, M., Yalcin, I., 2015. The anterior cingulate cortex is a critical hub for pain-induced depression. Biol Psychiatry 77, 236–245.

Bennett, G. J., Xie, Y. K., 1988. A peripheral mononeuropathy in rat that produces disorders of pain sensation like those seen in man. Pain 33, 87–107.

Berrocoso, E., Ikeda, K., Sora, I., Uhl, G. R., Sánchez-Blázquez, P., Mico, J. A., 2013. Active behaviours produced by antidepressants and opioids in the mouse tail suspension test. Int J Neuropsychopharmacol 16, 151–162.

Berrocoso, E., Rojas-Corrales, M. O., Mico, J. A., 2006. Differential role of 5-HT1A and 5-HT1B receptors on the antinociceptive and antidepressant effect of tramadol in mice. Psychopharmacology (Berl) 188, 111–118.

Camarena-Delgado, C., Llorca-Torralba, M., Suárez-Pereira, I., Bravo, L., López-Martín, C., Garcia-Partida, J. A., Mico, J. A., Berrocoso, E., 2022. Nerve injury induces transient locus coeruleus activation over time: role of the locus coeruleus-dorsal reticular nucleus pathway. Pain 163, 943–954.

Cardenas, A., Papadogiannis, A., Dimitrov, E., 2021. The role of medial prefrontal cortex projections to locus ceruleus in mediating the sex differences in behavior in mice with inflammatory pain. FASEB J 35, e21747.

Catuneanu, A., Paylor, J. W., Winship, I., Colbourne, F., Kerr, B. J., 2019. Sex differences in central nervous system plasticity and pain in experimental autoimmune encephalomyelitis. Pain 160, 1037–1049.

Chandler, D. J., Jensen, P., McCall, J. G., Pickering, A. E., Schwarz, L. A., Totah, N. K., 2019. Redefining Noradrenergic Neuromodulation of Behavior: Impacts of a Modular Locus Coeruleus Architecture. J Neurosci 39, 8239–8249.

Colburn, R. W., Lubin, M. L., Stone, D. J., Wang, Y., Lawrence, D., D’Andrea, M. R., Brandt, M. R., Liu, Y., Flores, C. M., Qin, N., 2007. Attenuated cold sensitivity in TRPM8 null mice. Neuron 54, 379–386.

Curzon, P., Rustay, N. R., Browman, K. E., 2009. Cued and Contextual Fear Conditioning for Rodents. In: Buccafusco, J. J., (Ed), Methods of Behavior Analysis in Neuroscience, Boca Raton (FL).

de la Rosa, T., Llorca-Torralba, M., Martinez-Cortes, A., Romero-Lopez-Alberca, C., Berrocoso, E., 2024. A Systematic Review and Meta-Analysis of Anxiety- and Depressive-Like Behaviors in Rodent Models of Neuropathic Pain. Biol Psychiatry Glob Open Sci 4, 100388.

Emery, E. C., Young, G. T., Berrocoso, E. M., Chen, L., McNaughton, P. A., 2011. HCN2 ion channels play a central role in inflammatory and neuropathic pain. Science 333, 1462–1466.

Gureje, O., Von Korff, M., Kola, L., Demyttenaere, K., He, Y., Posada-Villa, J., Lepine, J. P., Angermeyer, M. C., Levinson, D., de Girolamo, G., Iwata, N., Karam, A., Guimaraes Borges, G. L., de Graaf, R., Browne, M. O., Stein, D. J., Haro, J. M., Bromet, E. J., Kessler, R. C., Alonso, J., 2008. The relation between multiple pains and mental disorders: results from the World Mental Health Surveys. Pain 135, 82–91.

Hargreaves, K., Dubner, R., Brown, F., Flores, C., Joris, J., 1988. A new and sensitive method for measuring thermal nociception in cutaneous hyperalgesia. Pain 32, 77–88.

INFRAFRONTIER Consortium, 2015. INFRAFRONTIER--providing mutant mouse resources as research tools for the international scientific community. Nucleic Acids Res 43, D1171–1175.

Joshi, N., Chandler, D., 2020. Sex and the noradrenergic system. Handb Clin Neurol 175, 167–176.

Kasai, S., Yamamoto, H., Kamegaya, E., Uhl, G. R., Sora, I., Watanabe, M., Ikeda, K., 2011. Quantitative Detection of µ Opioid Receptor: Western Blot Analyses Using µ Opioid Receptor Knockout Mice. Curr Neuropharmacol 9, 219–222.

LeGates, T. A., Kvarta, M. D., Thompson, S. M., 2019. Sex differences in antidepressant efficacy. Neuropsychopharmacology 44, 140–154.

Lerman, S. F., Rudich, Z., Brill, S., Shalev, H., Shahar, G., 2015. Longitudinal associations between depression, anxiety, pain, and pain-related disability in chronic pain patients. Psychosom Med 77, 333–341.

Li, J., Wei, Y., Zhou, J., Zou, H., Ma, L., Liu, C., Xiao, Z., Liu, X., Tan, X., Yu, T., Cao, S., 2022. Activation of locus coeruleus-spinal cord noradrenergic neurons alleviates neuropathic pain in mice via reducing neuroinflammation from astrocytes and microglia in spinal dorsal horn. J Neuroinflammation 19, 123.

Llorca-Torralba, M., Camarena-Delgado, C., Suárez-Pereira, I., Bravo, L., Mariscal, P., Garcia-Partida, J. A., López-Martín, C., Wei, H., Pertovaara, A., Mico, J. A., Berrocoso, E., 2022. Pain and depression comorbidity causes asymmetric plasticity in the locus coeruleus neurons. Brain 145, 154–167.

Llorca-Torralba, M., Pilar-Cuéllar, F., Bravo, L., Bruzos-Cidon, C., Torrecilla, M., Mico, J. A., Ugedo, L., Garro-Martínez, E., Berrocoso, E., 2019a. Opioid Activity in the Locus Coeruleus Is Modulated by Chronic Neuropathic Pain. Mol Neurobiol 56, 4135–4150.

Llorca-Torralba, M., Suárez-Pereira, I., Bravo, L., Camarena-Delgado, C., Garcia-Partida, J. A., Mico, J. A., Berrocoso, E., 2019b. Chemogenetic Silencing of the Locus Coeruleus-Basolateral Amygdala Pathway Abolishes Pain-Induced Anxiety and Enhanced Aversive Learning in Rats. Biol Psychiatry 85, 1021–1035.

Luque, J. M., de Blas, M. R., Segovia, S., Guillamón, A., 1992. Sexual dimorphism of the dopamine-beta-hydroxylase-immunoreactive neurons in the rat locus ceruleus. Brain Res Dev Brain Res 67, 211–215.

MacDowell, K. S., Sayd, A., García-Bueno, B., Caso, J. R., Madrigal, J. L. M., Leza, J. C., 2017. Effects of the antipsychotic paliperidone on stress-induced changes in the endocannabinoid system in rat prefrontal cortex. World J Biol Psychiatry 18, 457–470.

Mariscal, P., Bravo, L., Llorca-Torralba, M., Razquin, J., Miguelez, C., Suárez-Pereira, I., Berrocoso, E., 2023. Sexual differences in locus coeruleus neurons and related behavior in C57BL/6J mice. Biol Sex Differ 14, 64.

McCall, J. G., Siuda, E. R., Bhatti, D. L., Lawson, L. A., McElligott, Z. A., Stuber, G. D., Bruchas, M. R., 2017. Locus coeruleus to basolateral amygdala noradrenergic projections promote anxiety-like behavior. Elife 6.

Miguelez, C., Morin, S., Martinez, A., Goillandeau, M., Bezard, E., Bioulac, B., Baufreton, J., 2012. Altered pallido-pallidal synaptic transmission leads to aberrant firing of globus pallidus neurons in a rat model of Parkinson’s disease. J Physiol 590, 5861–5875.

Mulvey, B., Bhatti, D. L., Gyawali, S., Lake, A. M., Kriaucionis, S., Ford, C. P., Bruchas, M. R., Heintz, N., Dougherty, J. D., 2018. Molecular and Functional Sex Differences of Noradrenergic Neurons in the Mouse Locus Coeruleus. Cell Rep 23, 2225–2235.

Paxinos, G., Franklin, K. B. J., 2019. The Mouse Brain in Stereotaxic Coordinates. Elsevier.

Porsolt, R. D., Le Pichon, M., Jalfre, M., 1977. Depression: a new animal model sensitive to antidepressant treatments. Nature 266, 730–732.

Rivera, B., Moreno, C., Lavanderos, B., Hwang, J. Y., Fernández-Trillo, J., Park, K. S., Orio, P., Viana, F., Madrid, R., Pertusa, M., 2021. Constitutive Phosphorylation as a Key Regulator of TRPM8 Channel Function. J Neurosci 41, 8475–8493.

Suárez-Pereira, I., Carrión, Á., 2015. Updating stored memory requires adult hippocampal neurogenesis. Sci Rep 5, 13993.

Tsang, A., Von Korff, M., Lee, S., Alonso, J., Karam, E., Angermeyer, M. C., Borges, G. L., Bromet, E. J., Demytteneare, K., de Girolamo, G., de Graaf, R., Gureje, O., Lepine, J. P., Haro, J. M., Levinson, D., Oakley Browne, M. A., Posada-Villa, J., Seedat, S., Watanabe, M., 2008. Common chronic pain conditions in developed and developing countries: gender and age differences and comorbidity with depression-anxiety disorders. J Pain 9, 883–891.

Valentino, R. J., Bangasser, D. A., 2016. Sex-biased cellular signaling: molecular basis for sex differences in neuropsychiatric diseases. Dialogues Clin Neurosci 18, 385–393.

Williams, J. T., North, R. A., Tokimasa, T., 1988. Inward rectification of resting and opiate-activated potassium currents in rat locus coeruleus neurons. J Neurosci 8, 4299-4306.

Wurzman, R., Forcelli, P. A., Griffey, C. J., Kromer, L. F., 2015. Repetitive grooming and sensorimotor abnormalities in an ephrin-A knockout model for Autism Spectrum Disorders. Behav Brain Res 278, 115–128.

Wyrofsky, R. R., Reyes, B. A. S., Yu, D., Kirby, L. G., Van Bockstaele, E. J., 2018. Sex differences in the effect of cannabinoid type 1 receptor deletion on locus coeruleus-norepinephrine neurons and corticotropin releasing factor-mediated responses. Eur J Neurosci 48, 2118–2138.

Yalcin, I., Bohren, Y., Waltisperger, E., Sage-Ciocca, D., Yin, J. C., Freund-Mercier, M. J., Barrot, M., 2011. A time-dependent history of mood disorders in a murine model of neuropathic pain. Biol Psychiatry 70, 946–953.

