## Supplemental Tables for "Sex differences in the impact of nerve injury on Locus Coeruleus function and behaviour in mice"

**Supplementary Table 1. Behavioural outcomes associated with neuropathic pain in male and female C57BL/6J mice.** Comprehensive overview of the behavioural tests conducted on male and female C57BL/6J mice, illustrating the progression and impact of neuropathic pain, highlighting sex differences and the role of the inhibition of the noradrenergic LC system.

| **BEHAVIOUR** | | **MALE** | | | **FEMALE** | | |
|---|---|---|---|---|---|---|---|
| | | **CCI-2/3w** | **CCI-7/11w** | **CCI-7/11w LC inhibition** | **CCI-2/3w** | **CCI-7/11w** | **CCI-7/11w LC inhibition** |
| **PAIN** | **Von Frey test** | ↓ mechanichal threshold | ↓ mechanichal threshold | | ↓ mechanichal threshold | ↓ mechanichal threshold | |
| | **Acetone test** | ↑ cold response | ↑ cold response | | ↑ cold response | ↑ cold response | |
| | **Plantar test** | | ↓ paw withdrawal threshold | | | ↓ paw withdrawal threshold | |
| **DEPRESSION** | **FST** | 0 | ↑ immobility time | | 0 | ↑ immobility time | |
| | **TST** | 0 | ↑ immobility time | ↓ immobility time | 0 | ↑ immobility time | 0 |
| **ANXIETY & FEAR** | **OF test** | | ↓ time in center | ↑ time in center | | 0 | 0 |
| | **FC test** | | ↑ freezing in conditioning phase | ↓ freezing in conditioning phase | | 0 | 0 |
| | **Stress-induced anxiety (LPS)** | | ↑ thigmotaxis | | | ↑ thigmotaxis | |
| **COGNITION** | **NOR** | | ↓ discrimination index | | | ↓ discrimination index | |

*0 represents no phenotype; Grey shading means DREADDs-mediated LC inhibition in CCI-7/11w CNO versus CCI-7/11w Saline mice; CCI, Chronic Constriction Injury; 2/3w, 2-3 weeks post CCI-surgery; 7/11w, 7-11 weeks post CCI-surgery; CNO, Clozapine-N-oxide; FC, Fear Conditioning; FST, Forced Swimming Test; LPS, Lipopolysaccharide; NOR, Novel Object Recognition; OF, Open Field; TST, Tail Suspension Test*

### Supplementary Table 2. Summary of the sample size of animals used in the study.

| | 2/3w | | | | Total number |
|---|---|---|---|---|---|
| | ♂Sham | ♂CCI | ♀Sham | ♀CCI | |
| **Cohort #1** | 10 | 10 | 10 | 10 | 40 |
| **Cohort #3** | 10 | 10 | 10 | 10 | 40 |
| | **7/11w** | | | | **Total number** |
| | **♂Sham** | **♂CCI** | **♀Sham** | **♀CCI** | |
| **Cohort #2** | 10 | 10 | 10 | 10 | 40 |
| **Cohort #4** | 10 | 10 | 10 | 10 | 40 |
| **Cohort #5** | 10 | 10 | 10 | 10 | 40 |
| **Cohort #7** | 9 | 12 | 8 | 9 | 38 |
| **Cohort #8** | 13 | 10 | 11 | 10 | 44 |
| **Cohort #9** | 5 | 6 | 6 | 6 | 23 |
| | | | | **Total** | 305 |

| | 7/11w | | | | | | | | Total number |
|---|---|---|---|---|---|---|---|---|---|
| | ♂Sham-Veh | ♂CCI-Veh | ♂Sham-LPS | ♂CCI-LPS | ♀Sham-Veh | ♀CCI-Veh | ♀Sham-LPS | ♀CCI-LPS | |
| **Cohort #6** | 11 | 9 | 8 | 12 | 9 | 7 | 10 | 9 | 75 |

| | 7/11w | | | | Total number |
|---|---|---|---|---|---|
| | ♂CCI-Sal | ♂CCI-CNO | ♀CCI-Sal | ♀CCI-CNO | |
| **Cohort #10** | 11 | 11 | 9 | 10 | 41 |
| **Cohort #11** | 11 | 11 | 11 | 8 | 41 |
| | | | | **Total** | 82 |

*CCI, Chronic Constriction Injury; 2/3w, 2-3 weeks post CCI-surgery; 7/11w, 7-11 weeks post CCI-surgery; CNO, Clozapine-N-oxide; LPS, Lipopolysaccharide; Sal, Saline; Veh, Vehicle*
